# Common biological features of *Mycobacterium tuberculosis* MmpL3 inhibitors

**DOI:** 10.1101/2025.03.31.646319

**Authors:** Lauren Ames, Renee Allen, Helena I. Boshoff, Laura Cleghorn, Curtis Engelhart, Dirk Schnappinger, Tanya Parish

## Abstract

MmpL3 is a promising new target for antitubercular drugs, but the microbiological properties of MmpL3 inhibitors are not fully understood. We compared the activity and mode of action of 11 structurally diverse compound series that target MmpL3. We confirmed activity was via MmpL3 using strains with differential expression of MmpL3. MmpL3 inhibitors had potent activity against replicating *M. tuberculosis*, with increased activity against intramacrophage bacilli and were rapidly bactericidal. MmpL3 inhibition induced cell wall stress concomitantly with a boost in ATP levels in *M. tuberculosis*. Mutation in MmpL3 conferred resistance to all series at different levels. Molecules did not negatively impact membrane potential, pH homeostasis or induce reactive oxygen species and were inactive against starved bacilli. Our study revealed common features related to the chemical inhibition of MmpL3, enabling the identification of off-target effects and highlighting the potential of such compounds as future drug candidates.

## Introduction

Tuberculosis (TB) remains a leading cause of mortality due to a single infectious disease, causing 10.8 million active infections and 1.25 million deaths in 2023^1^. Current treatment regimens for TB infections are lengthy, requiring four antibiotics taken for four to six months. Drug-resistant infections, accounting for 400,000 infections in 2023^1^, require an extended treatment time of up to 24 months with second line drugs. Many of the existing TB drugs are associated with adverse effects including visual impairment, hepatoxicity, neuropathy and QTc prolongation^2–4^. To improve treatment outcomes, there is an urgent need for more effective drugs that shorten treatment times and with fewer adverse effects. Identifying compounds with new targets is key to this goal.

*Mycobacterium tuberculosis*, the causative agent of TB, has a thick cell wall vital for survival in the host environment. The cell wall consists of a highly hydrophobic layer of mycolic acids linked to arabinogalactan which form an impermeable barrier to many antibiotics and is vital for adaptation to the host environment^5^. Preventing *M. tuberculosis* from maintaining its cell wall by treating with cell wall synthesis inhibitors, isoniazid and ethambutol^6–8^, has already proven effective in current TB treatment regimens. As a result, there is great interest in the discovery of compounds with other cell wall targets, such as MmpL3. MmpL3 is the sole transporter of mycolic acids in *M. tuberculosis*. It uses the proton motive force (PMF) to transfer TMM (trehalose monomycolate) across the inner membrane to the periplasmic space where it is incorporated into the mycolic acid layer by the Ag85 complex^9–11^.

MmpL3 is an attractive drug target since it is unique to *Mycobacteria* sp., is required for survival in macrophages^11,12^, for virulence and persistence in murine models of infection^13^ and its inhibition results in rapid cell death^11,13–15^. Numerous chemical scaffolds targeting MmpL3 have been described but none have reached approval for clinical use^16^. SQ109 is the furthest in development and reached clinical trials^17–19^ but may not be suitable for clinical use and likely has additional targets^20–22^.

Studies into the activity and mode of action of MmpL3 inhibitors have been confounded by the off-target effects of some of the most well-studied candidates. For example, SQ109 and BM212 appear to disrupt the PMF^20,21^, while others do not. SQ109 is active against *M. tuberculosis* in low oxygen whereas AU1235 and THPP-2 are inactive^20^. In addition, the fact that molecules have been studied in different laboratories under different experimental conditions complicates a direct comparison of activity. Understanding the biology of MmpL3 inhibitors is important in considering how drugs targeting this protein could be used in a multi-drug treatment regimen.

To understand the microbiological properties of MmpL3 inhibitors, we collected a set of structurally diverse compounds that target MmpL3 and simultaneously compared their activity against *M. tuberculosis*, their mechanism of action and interaction with MmpL3 mutant strains. Our study revealed a common set of features across MmpL3 inhibitors, some of which make these compounds promising drug candidates.

## Methods

### *M. tuberculosis* culture

Unless otherwise stated, experiments were performed using *M. tuberculosis* H37Rv-LP (ATCC 25618). *M. tuberculosis* strains were cultured in Middlebrook 7H9 medium plus 10 % v/v Middlebrook OADC supplement and 0.05 % w/v Tween 80 and incubated at 37 °C. For starvation, log phase bacteria were harvested and resuspended in PBS plus 0.05 w/v % Tyloxapol and incubated for 7d.

### Determination of minimum inhibitory concentration (MIC)

*M. tuberculosis* was grown to log phase and inoculated to a final OD_590_ of 0.02 in 96-well plates containing compounds as two-fold dilutions. Plates were incubated for 5 days at 37 °C. OD_590_ was read using a Synergy H4 plate reader. Dose response curves were fit using the variable Hill slope model from which IC_50_ and IC_90_ values, the concentration at which 50 % or 90 % growth inhibition occurred respectively, were determined.

### Determination of activity using recombinant strains

The assay was performed as previously described^23^. Briefly, the *mmpL3-TetON* strain was grown in 7H9 medium supplemented with 25 µg/mL kanamycin, 25 µg/mL zeocin +/- 500 ng/mL anhydrotetracycline (ATc). Cultures were diluted to OD_580_ of 0.01 in medium +/- ATc and 50 µL was dispensed into 384-well plates containing compound. Plates were incubated at 37 °C with 5 % CO_2._ OD_580_ was read after 14 d incubation.

### Determination of activity against intracellular bacilli

THP-1 cells were obtained from ATCC (TIB-202) and maintained in RPMI-1640 medium with 10 % FBS and incubated at 37 °C, 5 % CO_2_. THP-1 cells were differentiated by treatment with 80 nM PMA for 24 hours prior to overnight infection with an M. *tuberculosis* H37Rv-LP strain constitutively expressing luciferase from the pMV306hsp+LuxG13 plasmid^24^ at a multiplicity of infection (MOI) of 1:1. Infected cells were harvested using Accumax™ solution, resuspended in fresh cRPMI and inoculated to a final density of 9 x10^5^ cells/mL in 96-well plates containing compounds. Plates were incubated at 37 °C and 5 % CO_2_ for 72 hours. Luminescence was read using a Synergy H4 plate reader. Dose response curves were fit using the variable Hill slope model and the IC_50_ was determined.

### Determination of minimum bactericidal concentration (MBC)

*M. tuberculosis* H37Rv-LP was grown to log phase and inoculated to a final OD_590_ of 0.02 in 96-well plates containing compounds as two-fold dilutions. Plates were incubated at 37 °C. Samples were taken 0, 7 and 14 days, spotted onto 7H10 agar plates, and incubated at 37 °C for 2 weeks. The MBC (minimum bactericidal concentration) was determined visually as the lowest concentration at which no growth was observed.

### Determination of intracellular ATP levels

*M. tuberculosis* H37Rv-LP was grown to late log phase and inoculated at OD_590_ 0.04 in 96-well plates containing compounds as two-fold dilutions. Plates were incubated at 37 °C for 24 hours. ATP was quantified by adding 50 µL BacTiter-Glo™ reagent per well, incubating for 10 min and reading luminescence using a Synergy H4 plate reader. Kanamycin and Q203 were used as negative and positive controls, respectively.

### Induction of cell wall stress

The P_iniB_ reporter assay was adapted from Alland *et al.* 2000^25^. The *M. tuberculosis* P_iniB_-Lux strain, in which luciferase expression is controlled by the *iniB* promoter^26^, was grown in GAST-Fe medium and inoculated to a final OD_590_ of 0.02 in 96-well plates containing compounds as two-fold dilutions. Plates were incubated at 37 °C for 72 hours. 100 µL of substrate solution (28 µg/mL D-luciferin, 50 mM DTT in 1 M HEPES buffer) was added to each well and plates were incubated for 25 min at RT. Luminescence was read using a Synergy H4 plate reader. Ethambutol was used as a positive control.

### Induction of reactive oxygen species (ROS)

ROS was measured as previously described^27^. Briefly, *M. tuberculosis* H37Rv-LP was grown to an OD_590_ of 1, resuspended in 7H9-Tw + 40 µM 2’,7’-dichlorodihydrofluorescein diacetate (DCFDA) and incubated at 37 °C for 30 min. Cells were washed, resuspended in fresh 7H9-Tw medium and inoculated to a final OD_590_ of 0.5 in 96-well plates containing compound. Plates were incubated at 37 °C for 90 mins and fluorescence at Ex485/Em535 nm was measured using a Synergy H4 plate reader. Econazole was used as a positive control.

### Determination of membrane potential

*M. tuberculosis* H37Rv-LP was grown to late log phase and adjusted to an OD_590_ of 1 in fresh medium. 3,3’-diethyloxacarbocyanine iodide (DiOC_2_) was added to a final concentration of 15 µM and incubated at RT for 20 min. Cells were harvested, resuspended in fresh medium and inoculated to a final OD_590_ of 0.5 in 96-well plates containing compounds. Plates were incubated at 37 °C for 30 min. Fluorescence was read using an H4 Synergy plate reader at Ex488/Em530 and Ex488/Em610 nm. Carbonyl cyanide m-chlorophenyl hydrazone (CCCP) was used as a positive control.

### Determination of intracellular pH homeostasis

Intracellular pH was measured as previously described^28^. Briefly, *M. tuberculosis* H37Rv-LP expressing a pH-responsive ratiometric GFP (rGFP) from the pHLUOR plasmid^29^ was grown to late log phase, washed and resuspended in phosphocitrate buffer pH 4.5 (0.0896 M Na_2_HPO_4_, 0.0552 M citric acid, 0.05 % tyloxapol). The culture was inoculated to a final OD_590_ of 0.3 into 96-well assay plates containing compounds in two-fold dilutions. Plates were incubated at 37 °C for 48 hours. Fluorescence was measured using an H4 Synergy plate reader at Ex395/Em510 nm and Ex475/Em510 nm. Monensin was used as a control. A standard curve was generated using cell-free extracts as follows: *M. tuberculosis* H37Rv-LP rGFP was lysed using a bead beater then passed through a 0.2 micron filter to obtain cell-free extracts. Cell-free extracts (10 µg/mL protein) were adjusted to pH 5.5 to 8.5 and fluorescence measured to generate a ratio: the standard curve was fit using a four-parameter logistic model.

## Results

### Compound selection

The goal of our study was to understand the microbiological effects of chemical MmpL3 inhibition in *M. tuberculosis*. To avoid identifying effects related to specific chemical scaffolds, we included 11 chemically diverse compound series in our study (Figure 1). All 11 series originated from phenotypic hits identified in whole cell *M. tuberculosis* screens of chemical libraries (unpublished data) except for the pyrazole series which were derived from the previously described BM212 scaffold to improve its drug-like properties^30^. The selected series encompass varying stages of drug discovery in which activity against *M. tuberculosis* and drug-like properties are optimized. The DDU01, DDU02, DDU03, DDU04, and DDU05 series were early in discovery (hit assessment) in which some structure-activity-relationship exploration had been performed. The indolecarboxamide (ICA) series were the furthest progressed (pre-clinical candidate declaration) of the 11 series; activity and drug-like properties have been well-optimized, and compounds have proven efficacy in mouse models of infection^31^. The remaining series, pyrazole, ARI, 7-aza, oxazole, thiazole, were in mid-stage discovery (hit-to-lead or lead optimization). Two to three compounds representing each of the 11 series were selected for our study (Figure 1; compounds **1-32**). AU1235 was included as a reference since it is a well characterized MmpL3 inhibitor with no known secondary targets^10,14^.

**Figure 1:**
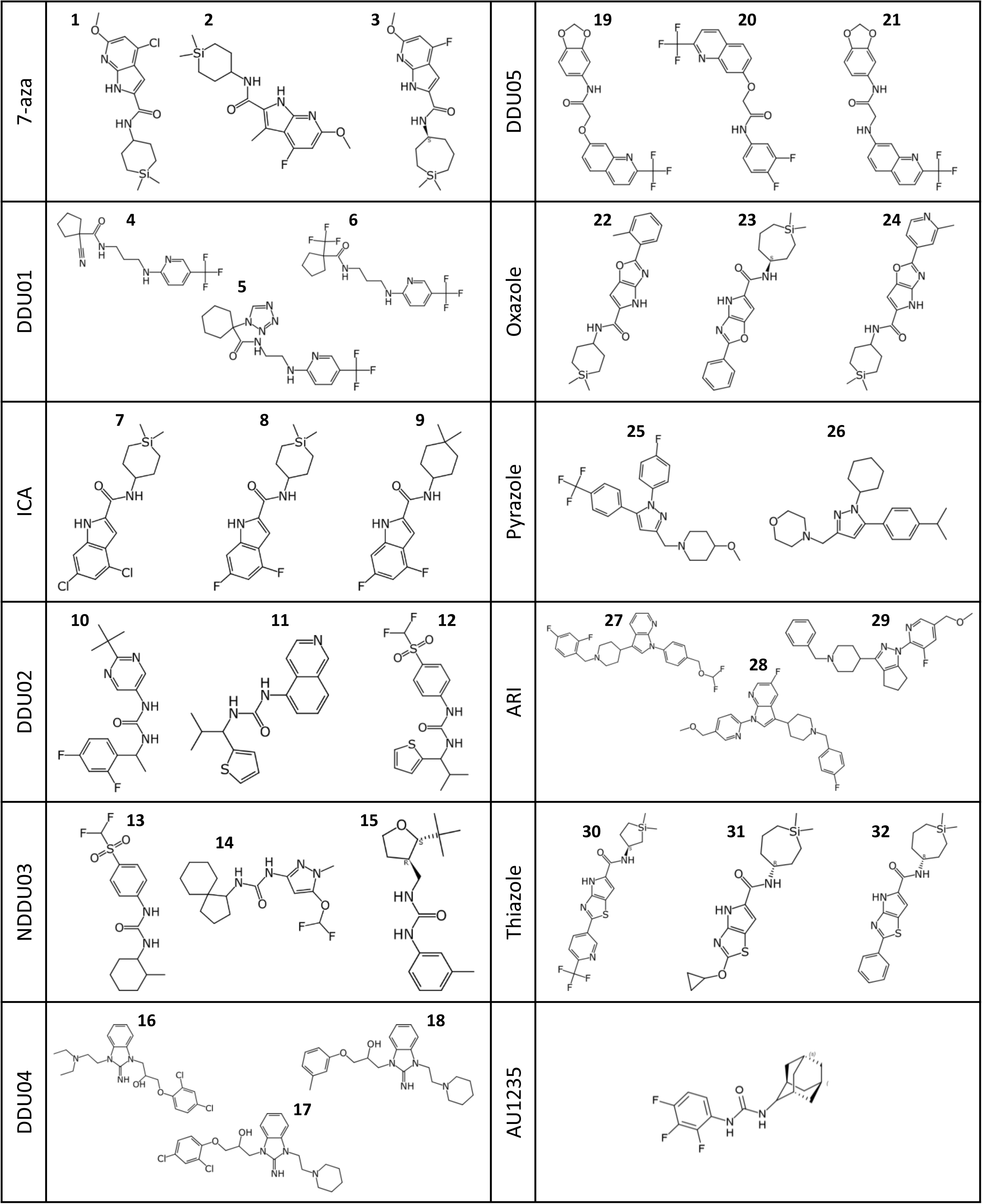
Chemical structures of MmpL3 inhibitors included in the study. Molecules are grouped by compound series.

### Compound series target MmpL3

To confirm MmpL3 as the target, compound activity was measured in *M. tuberculosis* strains with altered MmpL3 expression. The *mmpL3-TetON* strain was used in which MmpL3 expression is regulated by anhydrotetracycline^23^ (ATc). In the absence of ATc, expression of MmpL3 protein is decreased ∼80% and the *mmpL3-TetON* strain is more susceptible to MmpL3 inhibitors such as AU1235^23^ (Figure 2). Similarly, compounds across all 11 series were two- to six-fold more active against the *mmpL3-TetON* strain in the absence of ATc relative to the wild-type H37Rv (Figure 2). In the presence of ATc, MmpL3 expression in the *mmpL3-TetON* strain is increased by more than 7-fold^23^. Interestingly, MmpL3 overexpression conferred resistance to only a subset of the MmpL3 inhibitor compounds in our study and did not confer resistance to AU1235, a well-established MmpL3 inhibitor (Figure 2). Overall, altered activity against strains with differential MmpL3 expression is consistent with MmpL3 as the target of our compound series. Ethambutol was used as a negative control since MmpL3 under expression does not affect sensitivity^23^ (Figure 2).

**Figure 2:**
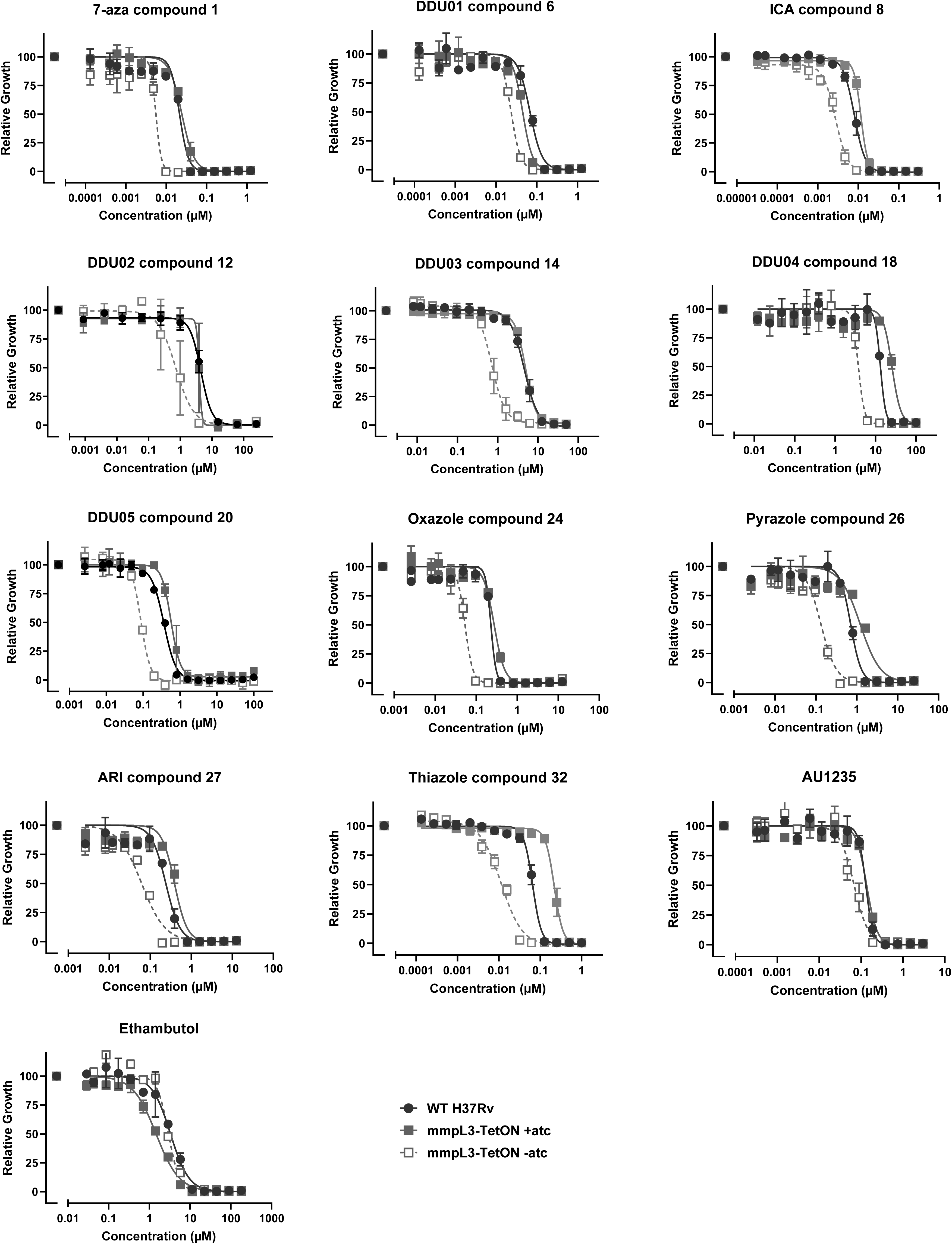
Differential expression of MmpL3 alters susceptibility to compounds. H37Rv (black circles) and the *mmpL3-TetON* strain with ATc (grey filled squares) or without ATc (grey empty squares) was exposed to compound for up to 14 days. OD_580_ was read. Plots shows mean and standard error for three independent experiments.

### MmpL3 mutations confer resistance

Mutations in the *M. tuberculosis mmpL3* gene confer resistance to a broad range of MmpL3 inhibitors^14,32–34^. To further confirm the target, we looked for resistance using an *M. tuberculosis* MmpL3 mutant strain, RM301 (MmpL3_F255L,V646M,F644I_). This strain was generated via multiple rounds of selection for resistance to different MmpL3 inhibitors and has been shown to confer resistance to structurally diverse MmpL3 inhibitor scaffolds^34^. The RM301 strain was >4-fold more resistant to most of our compounds relative to wild-type H37Rv-LP (25 out of 32 compounds; Table 2), consistent with MmpL3 being the primary target. RM301 was most resistant to the 7-aza series; activity against compound **1** was reduced >1639-fold relative to wild-type (Table 2). Of note, the RM301 strain conferred varying levels of resistance between the series, suggesting different interactions or binding affinities with MmpL3 (Table 2). However, mutations in the RM301 strain did not confer resistance to pyrazole or DDU05 series compounds, therefore we selected compound **26** and tested its activity against another MmpL3 mutant strain, MmpL3_S591I._ This strain was 23-fold more resistant to compound **26** relative to wild-type (Table 2) suggesting the pyrazoles may have a different mode of MmpL3 binding to the other series.

### MmpL3 inhibitors inhibit growth of *M. tuberculosis*

We measured the MICs (minimum inhibitory concentration) of our compounds against *M. tuberculosis* H37Rv-LP after five days in standard growth medium. All compounds inhibited growth of *M. tuberculosis*, some with excellent potency (IC_90_ < 1 µM) (Table 1). Compounds **7**, **8** and **9** from the ICA series were most active with an IC_90_ of 0.024, 0.029 and 0.071 µM respectively (Table 1). Compounds from the DDU02, DDU03 and DDU04 series were the least potent; compound **10** had the highest IC_90_ of 32 µM (Table 1). In general, compound activity correlated with stage in discovery as compounds in later stages of discovery were more potent than those in early discovery.

**Table 1.** Activity of compounds against *M. tuberculosis*. Inhibition of *M. tuberculosis* H37Rv-LP growth was measured after 5 days compound exposure in axenic culture (extracellular) or infected THP-1 cells (intramacrophage). Bactericidal activity of compounds against *M. tuberculosis* H37Rv-LP under replicating conditions (medium) and non-replicating conditions (starvation) after 14 days exposure. Values show the geometric mean of at least two independent experiments to 2 s.f.

| Series | Compound | Extracellular | | Intramacrophage | MBC ($\mu\text{M}$) | |
| --- | --- | --- | --- | --- | --- | --- |
| | | IC<sub>90</sub> ($\mu\text{M}$) | IC<sub>50</sub> ($\mu\text{M}$) | IC<sub>50</sub> ($\mu\text{M}$) | Medium | Starvation |
| 7-aza | 1 | 0.061 | 0.050 | 0.014 | <0.2 | 100 |
|  | 2 | 0.14 | 0.11 | 0.024 | <0.39 | >100 |
|  | 3 | 0.17 | 0.061 | 0.022 | <0.2 | 25 |
| DDU01 | 4 | 2.0 | 1.2 | 0.62 | 2.4 | >100 |
|  | 5 | 5.5 | 4.3 | 6.2 | 9.4 | >100 |
|  | 6 | 0.19 | 0.094 | 0.057 | <0.39 | >100 |
| ICA | 7 | 0.024 | 0.020 | 0.016 | <0.2 | >100 |
|  | 8 | 0.029 | 0.017 | 0.0060 | <0.2 | >100 |
|  | 9 | 0.071 | 0.038 | 0.0094 | <0.2 | >75 |
| DDU02 | 10 | 32 | 10 | 6.0 | 25 | >100 |
|  | 11 | 8.6 | 6.6 | 4.5 | 13 | >100 |
|  | 12 | 13 | 5.7 | 1.2 | 9.4 | 100 |
| DDU03 | 13 | 11 | 9.5 | 2.0 | 13 | 75 |
|  | 14 | 8.3 | 4.2 | 3.2 | 7.8 | >100 |
|  | 15 | 20 | 9.2 | 5.7 | 27 | >100 |
| DDU04 | 16 | 12 | 6.0 | 3.4 | 6.3 | 25 |
|  | 17 | 3.0 | 1.9 | 2.5 | 3.1 | 50 |
|  | 18 | 15 | 7.6 | 4.8 | 13 | >100 |
| DDU05 | 19 | 8.9 | 4.1 | 12 | 38 | >100 |
|  | 20 | 6.1 | 2.1 | 1.9 | 9.4 | >100 |
|  | 21 | 4.8 | 1.3 | 3.9 | >100 | >100 |
| Oxazole | 22 | 0.48 | 0.31 | 0.060 | <0.39 | >100 |
|  | 23 | 0.078 | 0.061 | 0.017 | <0.2 | >100 |
|  | 24 | 0.62 | 0.46 | 0.062 | 1.2 | >75 |
| Pyrazole | 25 | 0.27 | 0.12 | 0.024 | <0.39 | 100 |
|  | 26 | 2.3 | 1.1 | 0.27 | 1.6 | >100 |
| ARI | 27 | 0.58 | 0.31 | 0.063 | 0.78 | >100 |
|  | 28 | 1.1 | 0.65 | 0.24 | 1.6 | >100 |
|  | 29 | 2.3 | 1.0 | 0.19 | 3.1 | >100 |
| Thiazole | 30 | 0.18 | 0.044 | 0.019 | <0.2 | >100 |
|  | 31 | 0.19 | 0.081 | 0.051 | <0.2 | >100 |
|  | 32 | 0.088 | 0.053 | 0.0057 | <0.2 | >100 |
| AU1235 | - | 0.22 | 0.13 | 0.063 | 0.59 | >100 |

**Table 2.**
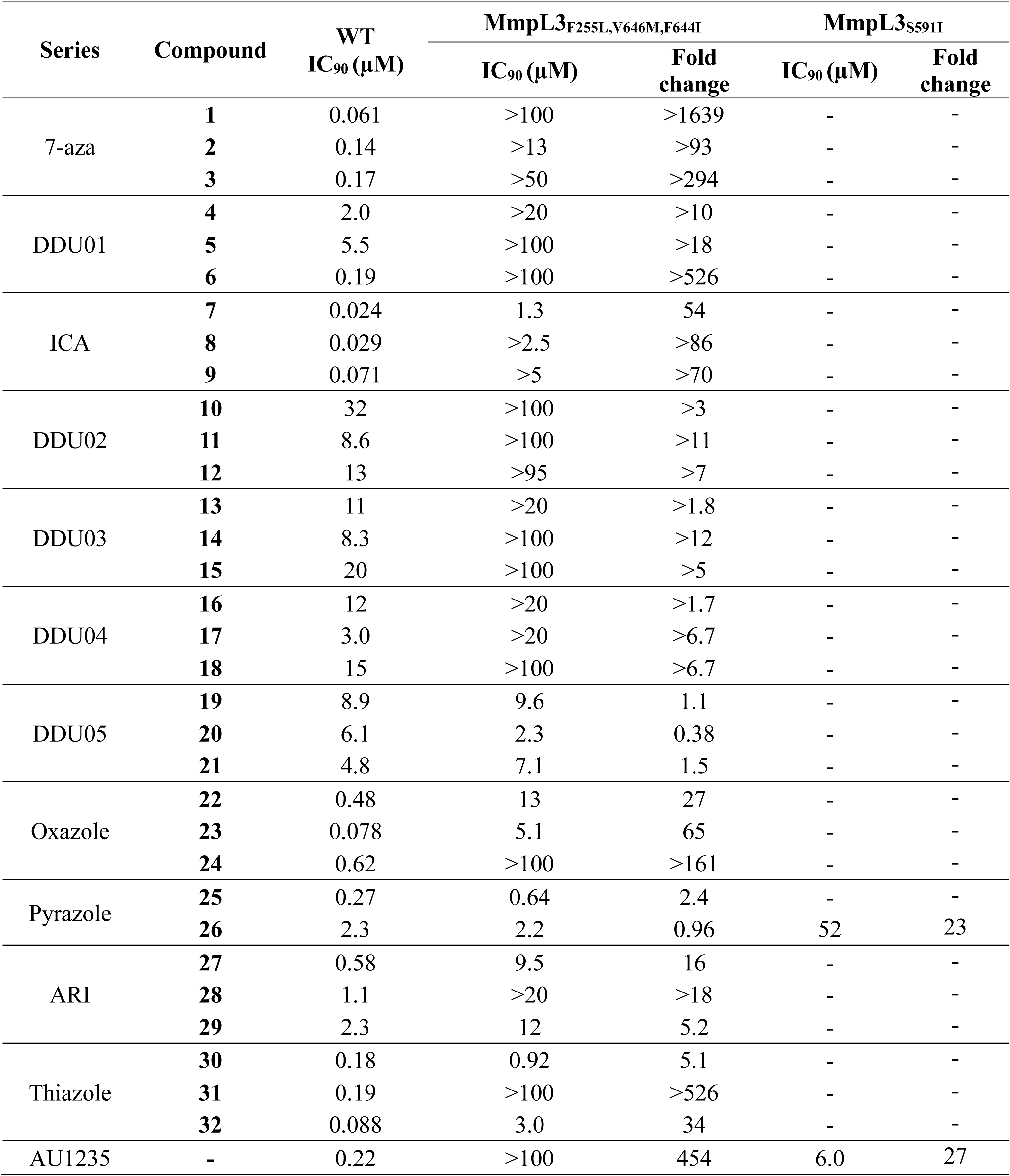
Compound activity against *M. tuberculosis* MmpL3 mutant strains. Inhibition of growth was measured after 5 days exposure to compound. IC_90_ values show the geometric mean of at least two independent experiments to 2 s.f. Fold change was calculated relative to WT IC_90_.

### MmpL3 inhibitors have enhanced activity against intracellular bacteria

We measured the activity of compounds against intramacrophage *M. tuberculosis.* Differentiated THP-1 cells were infected with *M. tuberculosis* H37Rv-LP and exposed to compound for three days. We found that all compounds across all 11 series, as well as AU1235, strongly inhibited growth of intramacrophage *M. tuberculosis* (Table 1). As with extracellular activity, compound activity correlated with stage in discovery. The ICA series was most potent; the intramacrophage IC_50_ for compounds **7**, **8** and **9** was 0.063, 0.0060 and 0.0094 µM respectively (Table 1). Compound **19** from the DDU05 series was least potent with an intramacrophage IC_50_ of 12 µM (Table 1). Comparison of intramacrophage and extracellular activity revealed that most compounds, including AU1235, were up to 10-fold more active against intramacrophage *M. tuberculosis* (Figure 3). The increased potency against intramacrophage *M. tuberculosis* was observed across chemically distinct compound series so is likely a feature of MmpL3 inhibition rather than an effect related to a particular chemical structure.

**Figure 3:**
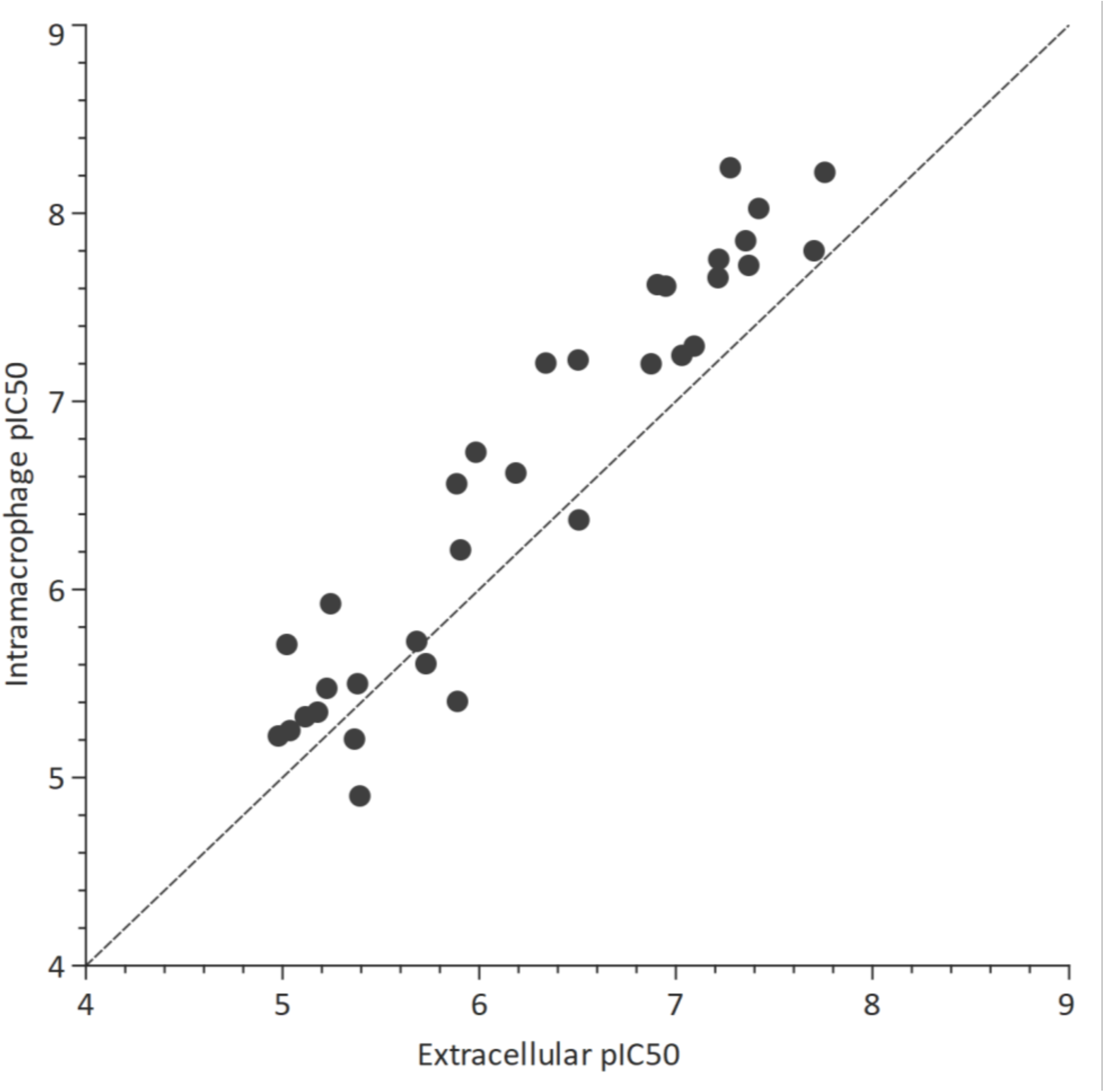
Activity of compounds against *M. tuberculosis* H37Rv-LP in axenic culture (extracellular) and infected THP-1 cells (intramacrophage). Points represent the geometric mean of at least two independent replicates. Dotted line shows *y* = *x*.

### MmpL3 inhibitors are bactericidal against replicating *M. tuberculosis*

We next compared the bactericidal activity of compounds. The minimum bactericidal concentration (MBC) was determined after 14 days exposure to compound in standard growth medium in which the bacilli are in a replicating state. Almost all compounds were bactericidal against *M. tuberculosis* in this condition, a number of which were bactericidal at sub-micromolar concentrations (Table 1). Compounds from the Thiazole and ICA series had most potent bactericidal activity with an MBC less than 0.2 µM (the lowest concentration tested) (Table 1). DDU05 series compound **21** was the only compound for which no bactericidal activity against replicating bacteria was observed up to 100 µM (the highest concentration tested) (Table 1). Across all series, the concentration required for bactericidal activity (MBC) was very close to the inhibitory concentration (IC_90_; Table 1). Such potent bactericidal activity is an unusual but promising property for antimicrobial compounds.

Since *M. tuberculosis* can also exist in a non-replicating state in the host, we looked for bactericidal activity against *M. tuberculosis* cells starved for seven days prior to compound exposure. All compounds, including AU1235, had little or no bactericidal activity against starved cells at the concentrations tested (up to 100 µM; Table 1). Thus, in contrast be previous studies using SQ109^20^, we found that MmpL3 inhibitors are not active against bacteria in a non-replicating state. This was not surprising since synthesis of the mycolic acid layer, and therefore mycolic acid export, is less important in cells that are not growing. However, the different methods used to induce a non-replicating state (low oxygen compared to nutrient starvation) between studies should be considered.

### MmpL3 inhibitors induce cell wall stress

To confirm the mode of action of MmpL3 inhibitors, we ran all compounds through a series of assays testing effects on different biological pathways. We first measured induction of cell wall stress using an *M. tuberculosis* reporter strain in which luciferase expression is controlled by the *iniB* promoter which is induced by cell wall stress^26^. As anticipated for inhibitors of mycolic acid transport, all compounds induced the P_iniB_ reporter indicating cell wall stress (Figure 4). Test compounds induced the reporter at similar levels to ethambutol, an inhibitor of cell wall arabinogalactan synthesis^7,8^. Reporter signal was strongest around the IC_50_ for each compound linking compound activity to the induction of cell wall stress.

**Figure 4:**
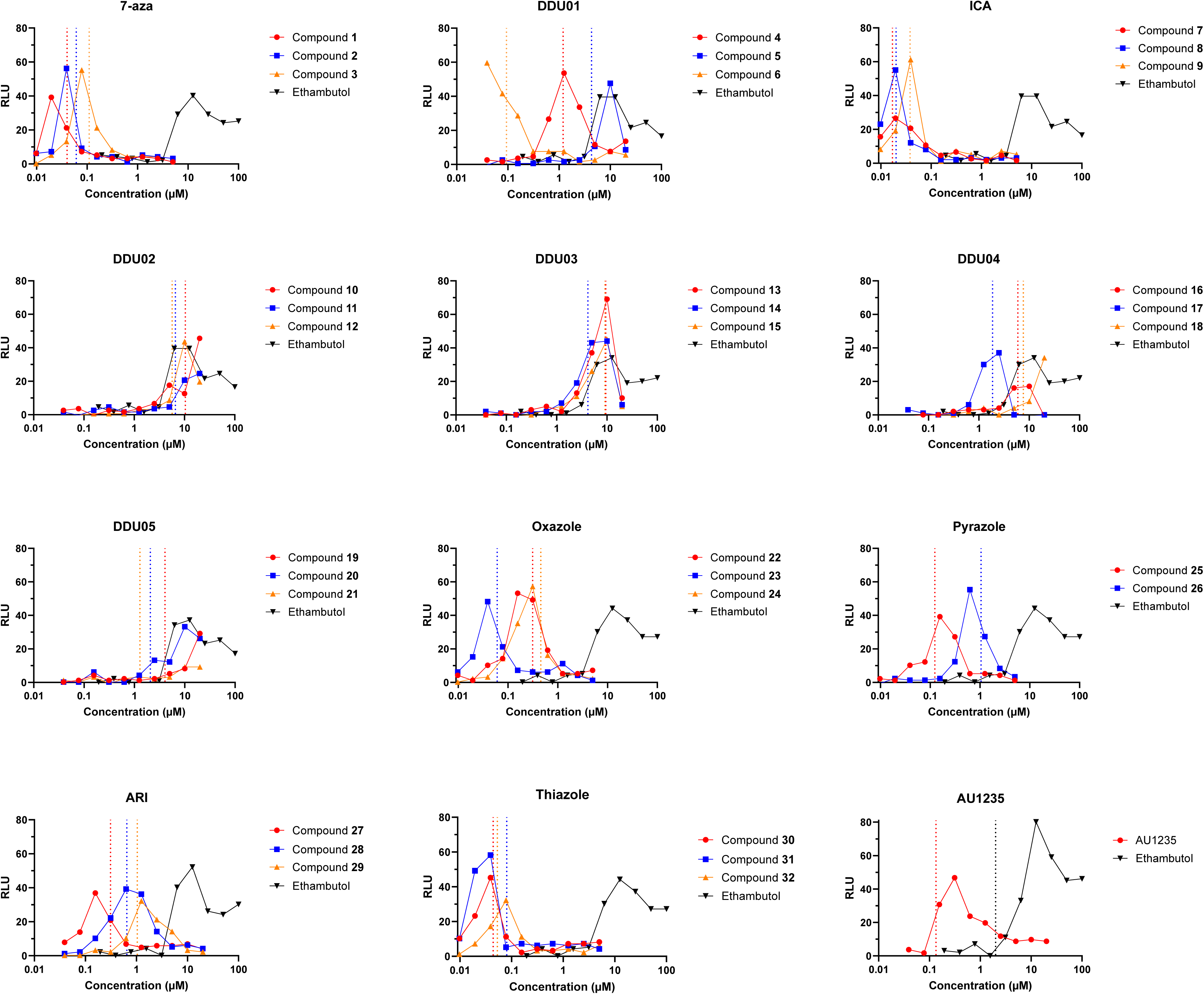
MmpL3 inhibitors induce cell wall stress. A *M. tuberculosis* strain expressing a luciferase operon under control of the cell wall stress-induced iniBAC promoter was exposed to compounds and luminescence (RLU) was measured. Ethambutol was used as a positive control. Plots show one of at least two independent experiments. Dotted vertical lines represent the IC_50_.

### MmpL3 inhibitors induce an ATP boost

We measured intracellular ATP following compound exposure. All compounds caused a substantial increase in ATP levels in our assay (Figure 5). This increase occurred at the concentration required to inhibit bacterial growth, suggesting that chemical MmpL3 inhibition induces an ATP burst in *M. tuberculosis.* Q203, a QcrB inhibitor^35^, was used as a control for ATP depletion and demonstrated the expected decrease in ATP levels at subinhibitory concentrations. Kanamycin (negative control) lead to a drop in ATP levels concomitant with decreased growth.

**Figure 5:**
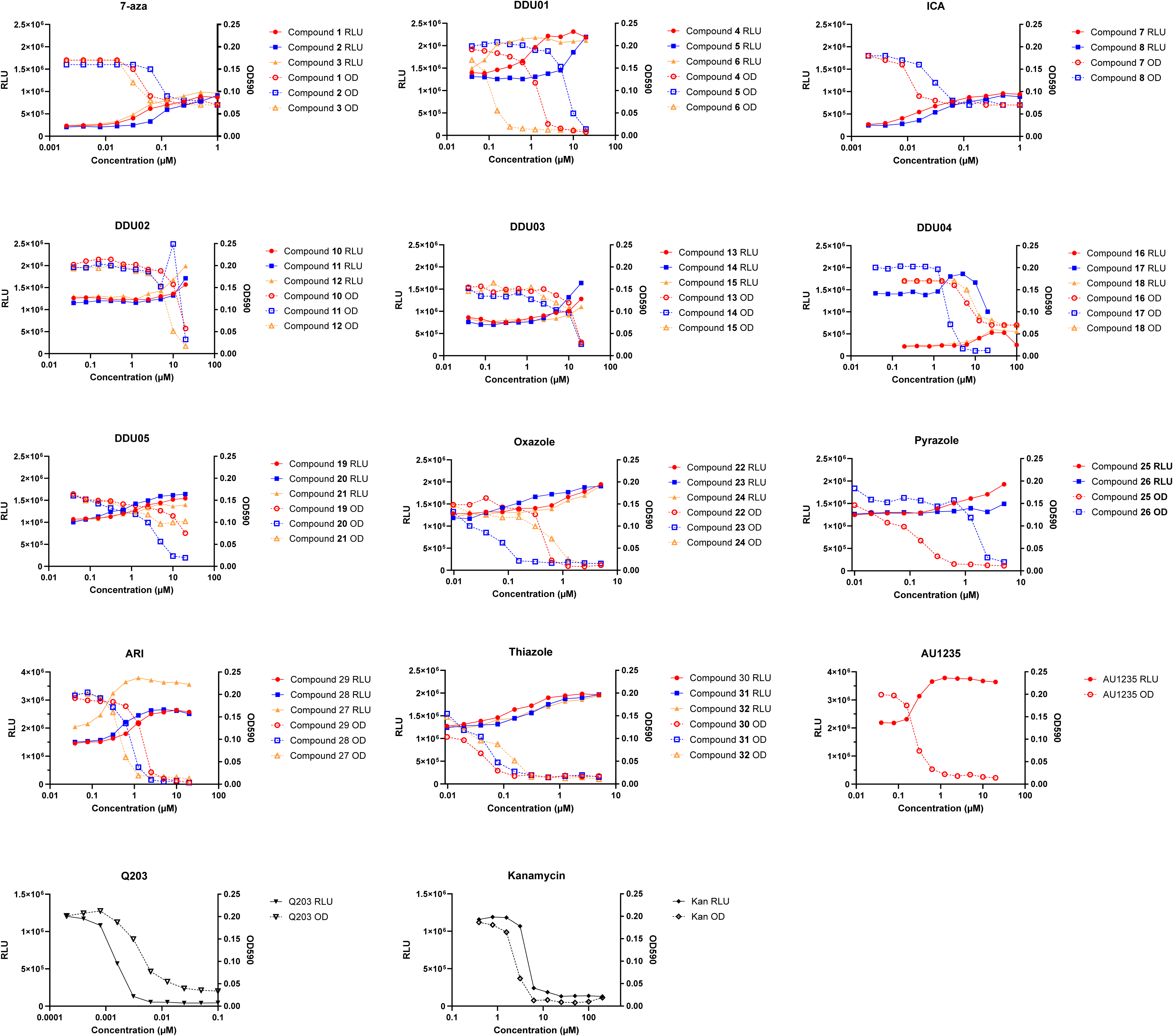
MmpL3 inhibitors induce an ATP boost. *M. tuberculosis* H37Rv-LP was exposed to compounds and ATP levels measured after 24 hours by addition of Bac-Titer Glo reagent and reading luminescence (RLU). OD_590_ was measured after 5 days. Q203 was used as a positive control (ATP depletion) and kanamycin was used as a negative control (no effect). Plot show one representative of at least two independent replicates.

This ATP boost in response to MmpL3 inhibition has not been previously described in *M. tuberculosis*. However, our finding is consistent with studies in other *Mycobacteria* sp., such as *Mycobacterium abscessus* and *Mycobacterium bovis* BCG, where treatment with cell wall inhibitors lead to increased ATP concentrations^36,37^, suggesting the ATP boost is a general response to cell wall perturbations. Further work is required to understand the mechanistic link between cell wall inhibition and increased ATP generation in *Mycobacteria* sp.

### MmpL3 inhibitors to do not affect ROS, membrane potential or pH homeostasis

Compounds were also tested for effects on reactive oxygen species (ROS) generation, dissipation of membrane potential and disruption of intracellular pH. ROS were quantified using 2′,7′-Dichlorofluorescin diacetate (DCFDA), an oxidation-sensitive fluorescent dye with econazole used as a positive control^27^.

Changes in membrane potential were detected using the fluorescent probe 3,3’-Diethyloxacarbocyanine iodide (DiOC2(3)) with monensin, an ionophore, as a positive control. The ability to maintain intracellular pH was tested using *M. tuberculosis* expressing a pH-dependent ratiometric GFP^28^. In general, none of the compounds induced ROS, or disrupted membrane potential or pH homeostasis (Figure 6; Figures S1 to S3). The exceptions were compound **24** from the oxazole series and compound **20** from the DDU05 series which induced intracellular alkalinization (Figure 6, Figure S2), suggesting these molecules have a secondary target. Our finding that structurally diverse MmpL3 inhibitors do not dissipate membrane potential suggest that the effect SQ109 has on membrane potential^20,21^ is structure specific, perhaps as a consequence of a second target rather than MmpL3 inhibition.

**Figure 6:**
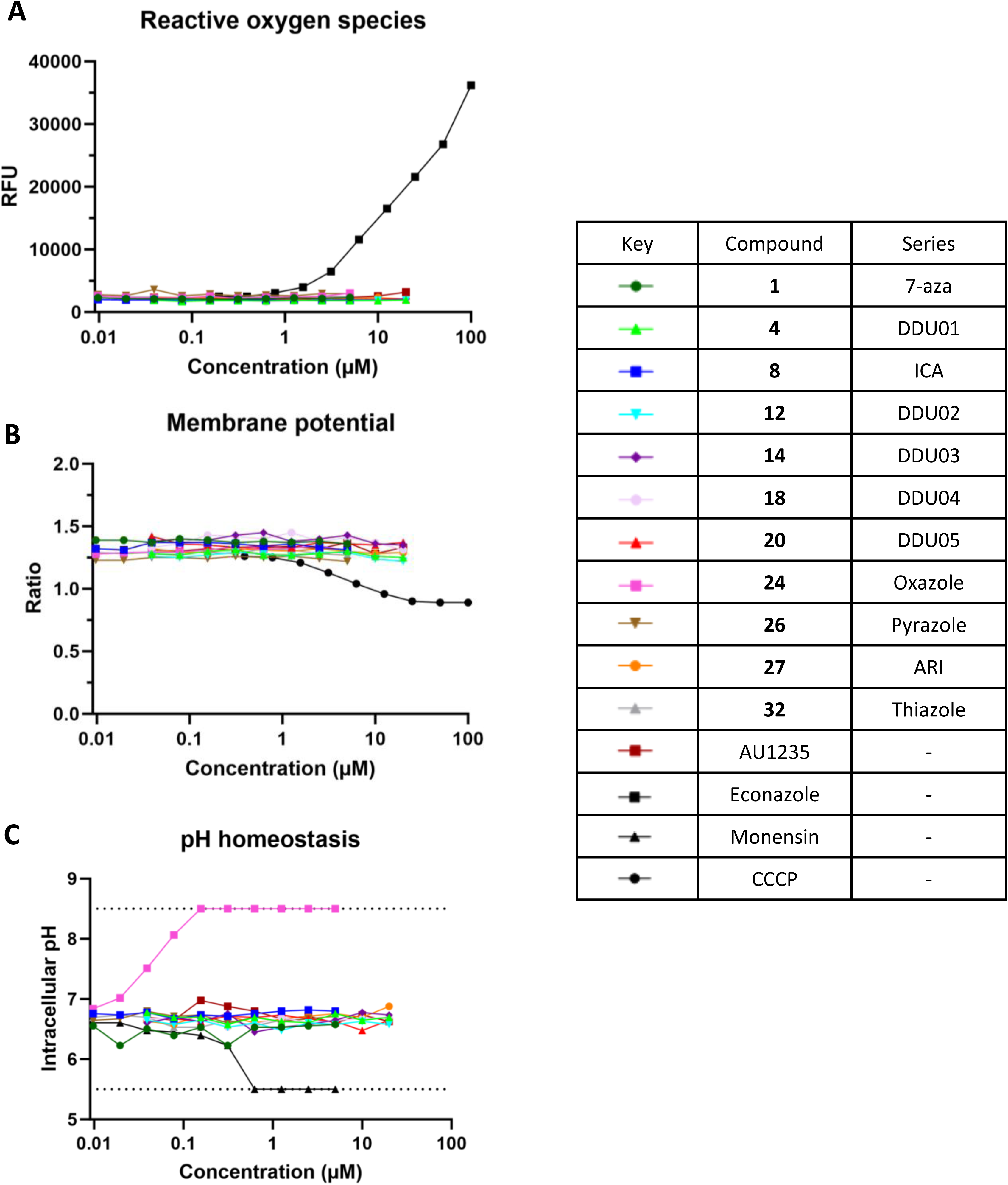
MmpL3 inhibitors do not induce reactive oxygen species (ROS) production, disrupt membrane potential or maintenance of intracellular pH. *M. tuberculosis* H37Rv-LP was exposed to compound. ROS was quantified using oxidation-sensitive fluorescent dye 2′,7′-Dichlorofluorescin diacetate (DCFDA) and measured fluorescence Ex485/Em535 nm (RFU). Membrane potential was measured using cells stained with fluorescent probe 3,3’-Diethyloxacarbocyanine iodide (DiOC2(3)) and calculating the ratio between Ex488/Em530 nm and Ex488/Em530 nm fluorescence reads. Intracellular pH was measured using an *M. tuberculosis* strain expressing a ratiometric GFP. Fluorescence was read at Ex390/Em505 nm and Ex475/Em510 nm and the ratio calculated. Intracellular pH was interpolated using a standard curve. Dotted lines represent the limit of intracellular pH measurement. Plots show one representative of at least two independent experiments.

Taken together, our assays confirmed the mode of action of MmpL3 inhibitors, which induced cell wall stress and increased ATP levels. Changes in membrane potential, pH homeostasis and ROS are not a general feature of MmpL3 inhibition, but rather off-target effects related to specific chemical scaffolds.

## Discussion

We performed a systematic comparison of 11 structurally diverse MmpL3 inhibitor series in which 32 representative compounds were run in parallel through a set of biological assays aimed at increasing our understanding of activity against *M. tuberculosis*, mode of action and interaction with MmpL3 mutant strains. AU1235 was chosen as a reference for the study since there is no evidence for off-target activity, to our knowledge.

We confirmed MmpL3 as the target of our compounds using MmpL3 mutant strains. Most MmpL3 inhibitors, including AU1235, SQ109 and NITD-349, bind the same pocket site and induce a conformation change in the MmpL3 protein structure that disrupts two Asp-Try pairs required for proton transport^38^. The same mutations that conferred resistance to AU1235 (MmpL3_F255L,V646M,F644I_ and MmpL3_S591I_), also conferred resistance to our compounds suggesting a similar mode of binding to MmpL3. The fact that MmpL3 mutants conferred different fold-change in resistance to each compound series could suggest different binding affinities to MmpL3. The specificity of resistance to certain compounds series in our study also suggests different modes of binding within the drug pocket. Further understanding the global landscape of MmpL3 mutations and how they relate to resistance will be key to predicting clinical resistance to MmpL3 inhibitors in future treatment regimens.

Compounds from all 11 series behaved similarly to AU1235, highlighting a set of common biological features of MmpL3 inhibitors: a) potent growth inhibition of *M. tuberculosis*; b) bactericidal activity against replicating bacilli; c) increased potency against intramacrophage *M. tuberculosis*; d) induction of cell wall stress; and e) an ATP boost. The identification of core biological features of MmpL3 inhibitors in this study has reinforced the potential of such compounds as potential antitubercular drugs. Many compounds had excellent bactericidal activity and inhibitory activity against intramacrophage *M. tuberculosis*. The contribution of these properties to treatment shortening warrants further study. It will also aid the identification of off-target activity in future drug discovery efforts and inform how such compounds may be used in combinatorial treatment regimens

## Acknowledgements

We thank Nikki Nguyen and Stephanie Jade Anover-Sombke for technical assistance.

---

**Figure S1:**
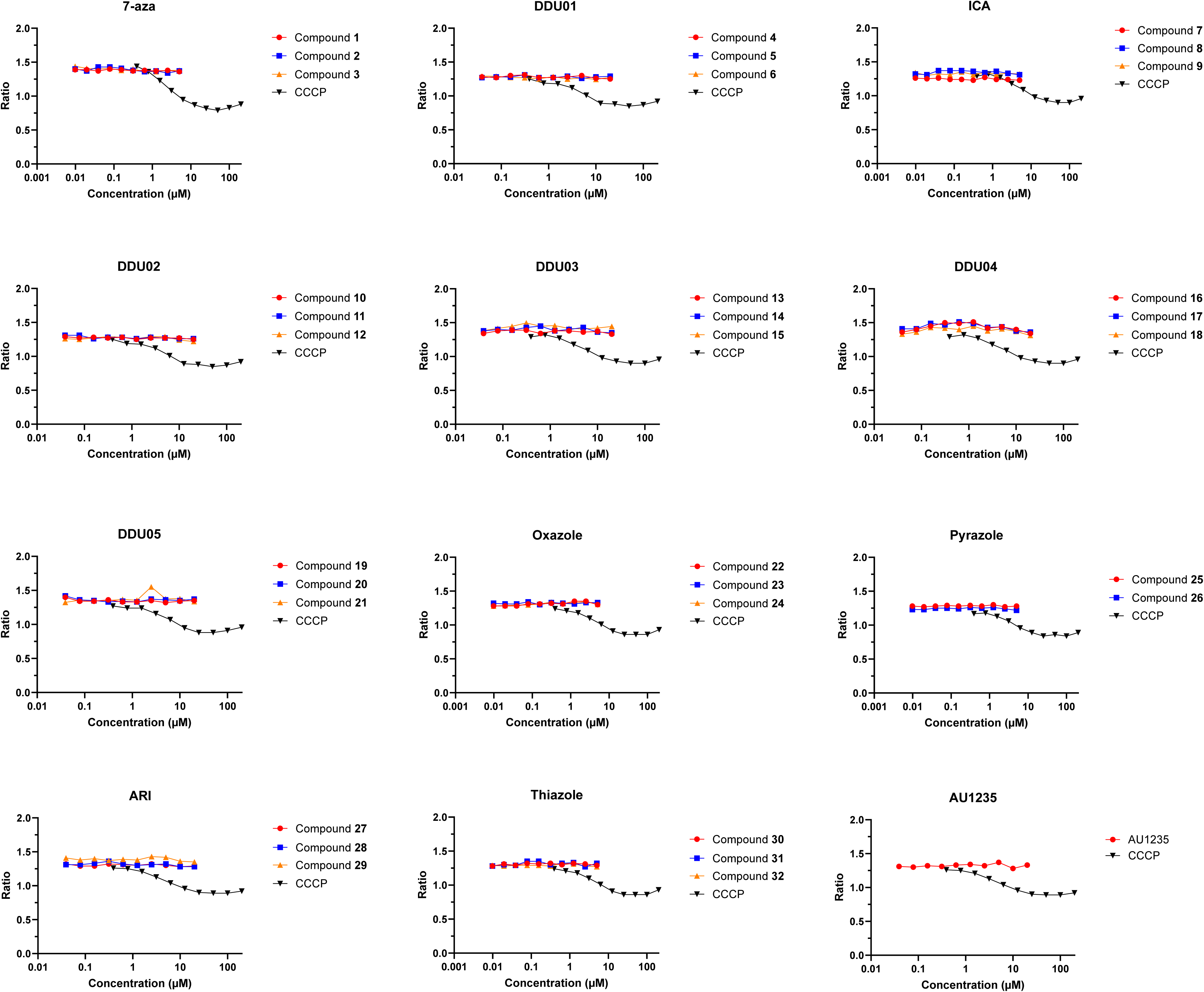
Compounds do not affect membrane potential. *M. tuberculosis* H37Rv-LP cells treated with DiOC2 were exposed to compound for 30 minutes and fluorescence was read (Ex488/Em530 nm and Ex488/Ex610 nm). The ratio between fluorescence reads was calculated. CCCP was used as a positive control. Plots show one representative of at least two independent experiments.

**Figure S2:**
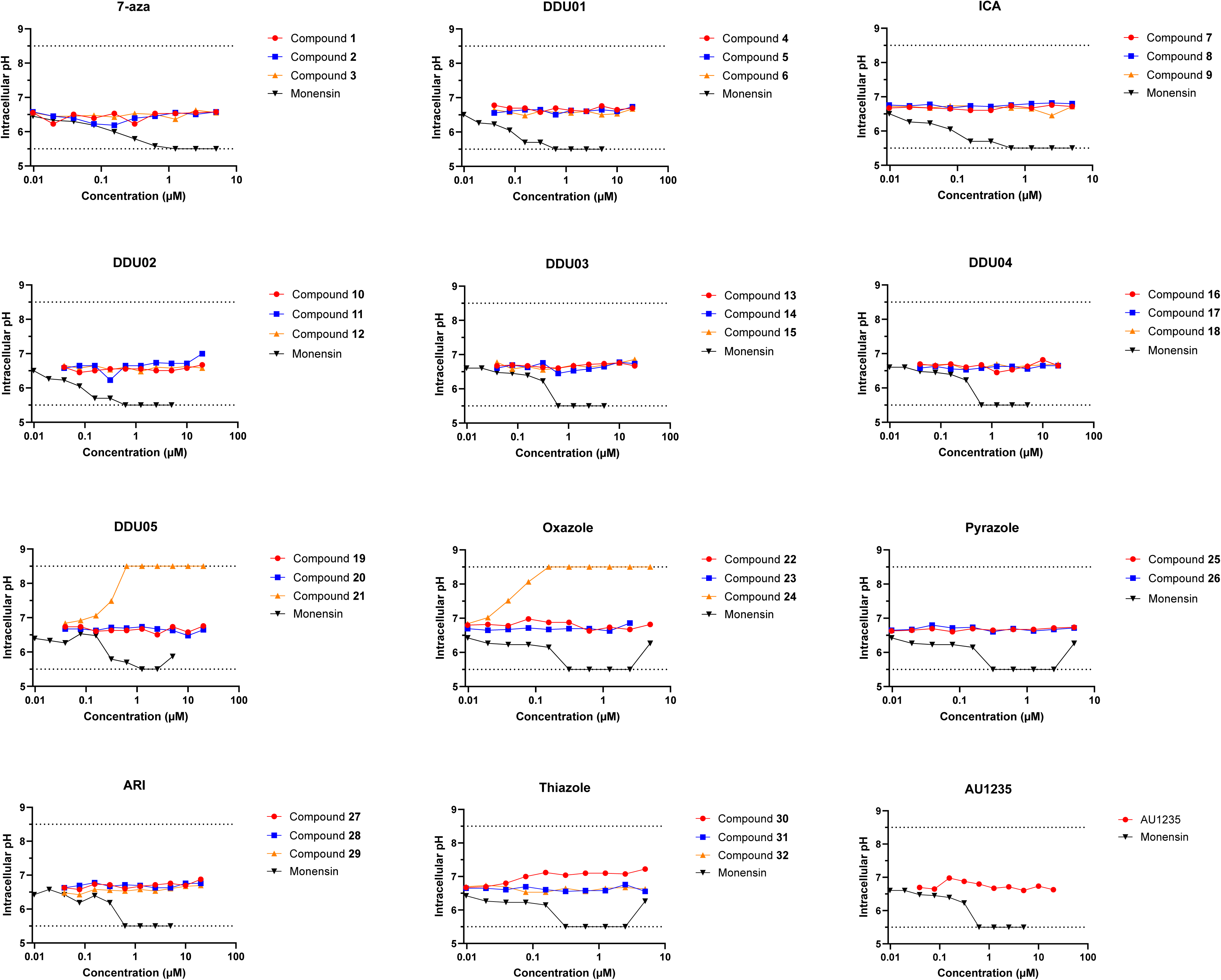
Compounds do not affect intracellular pH homeostasis. *M. tuberculosis* H37Rv-LP expressing a ratiometric GFP was exposed to compound in pH 4.5 buffer for 48 hours. Fluorescence was read (Ex395/Em510 nm and Ex475/Ex510 nm) and converted to pH using a standard curve. Monensin was used as a positive control. Plots show one representative of at least two independent experiments. Dotted lines show the measurable pH range in our assay.

**Figure S3:**
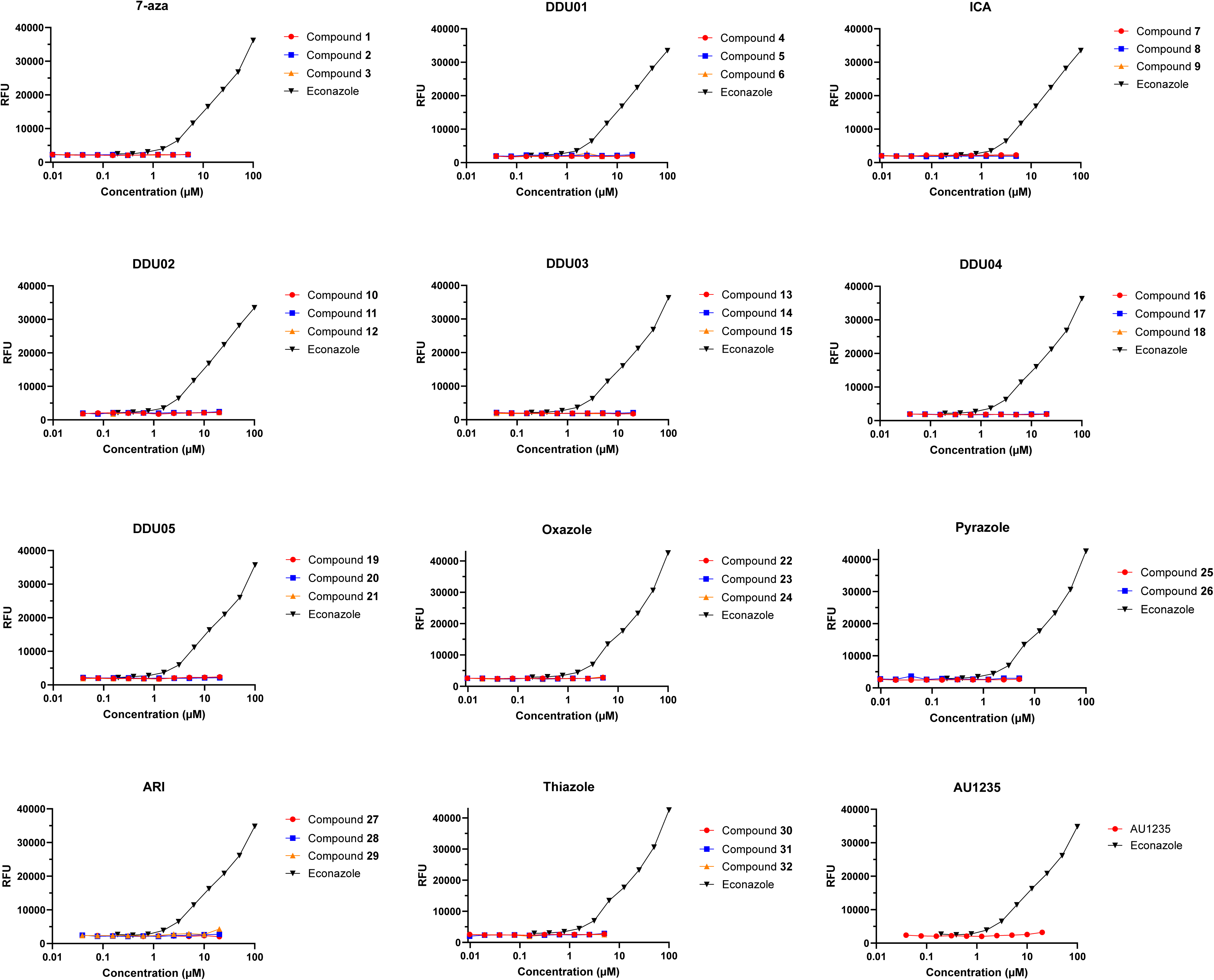
Compounds do not affect reactive oxygen species generation. *M. tuberculosis* H37Rv-LP cells treated with fluorogenic dye DCFA were exposed to compounds for 30 minutes and fluorescence was read (Ex485/Em535 nm). Econazole was used as a positive control. Plots show one representative of at least two independent experiments.

